# Encoding Variability: When Pattern Reactivation Does Not Benefit Context Memory

**DOI:** 10.1101/586446

**Authors:** Carolin Sievers, Fraser W. Smith, Janak Saada, Jon S. Simons, Louis Renoult

## Abstract

A growing body of evidence suggests that neural pattern reactivation supports successful memory formation across multiple study episodes. Previous studies investigating the beneficial effects of repeated encoding typically presented the same stimuli repeatedly under the same encoding task instructions. In contrast, repeating stimuli in different contexts is associated with superior item memory, but poorer memory for contextual features varying across repetitions. In the present functional magnetic-resonance imaging (fMRI) study, we predicted dissociable mechanisms to underlie the successful formation of context memory when the context in which stimuli are repeated is either held constant or varies at each stimulus presentation. Twenty participants studied names of famous people four times, either in the same task repeatedly, or in four different encoding tasks. This was followed by a surprise recognition memory test, including a source judgement about the encoding task. Behaviourally, different task encoding compared to same task encoding was associated with fewer correct context memory judgements but also better item memory, as reflected in fewer misses. Searchlight representational similarity analysis revealed fMRI pattern reactivation in the posterior cingulate cortex to be higher for correct compared to incorrect source memory judgements in the same task condition, with the opposite pattern being observed in the different task condition. It was concluded that higher levels of pattern reactivation in the posterior cingulate cortex index generalisation across context information, which in turn may improve item memory performance during encoding variability but at the cost of contextual features.

## Introduction

In our daily lives, we commonly encounter the same information multiple times; sometimes in different contexts or situations. It is such additional contextual information that is the essence of episodic memory and gives us the ability to re-experience personal past events (Schacter, Wagner, & Buckner, 2000; Tulving, 2002). While repetition typically improves our memory for core features of a stimulus, repeated encounters in different situations can interfere with our ability to remember contextual details that may vary across repetitions (e.g., Kim et al., 2012).

A large amount of existing research demonstrates that repeated memory encoding is associated with improved subsequent memory performance (e.g., Baddeley, 1978; Cepeda, Pashler, Vul, Wixted, & Rohrer, 2006; Crowder, 1976; Davachi, Maril, & Wagner, 2001; Glenberg, Smith, & Green, 1977; Greene, 1987; Mechanic, 1964; Opitz, 2010; Ranganath, Cohen, & Brozinsky, 2005; Reagh & Yassa, 2014; Van Strien, Hagenbeek, Stam, Rombouts, & Barkhof, 2005). However, the cognitive and neural mechanisms that might explain such repetition-related memory improvements are still highly contested. Two opposing theoretical frameworks are the reactivation view and the encoding variability view. According to the reactivation view, repeated encoding reactivates existing memory traces, thereby making the representation more stable and durable (Benjamin & Tullis, 2010; Thios & D’Agostino, 1976). On the other hand, the encoding variability view proposes that a stimulus is uniquely encoded at each of its presentations, which leads to an increase in number and variety of available retrieval cues (Bower, 1972; Hintzman, 1986; Martin, 1968; Nadel & Moscovitch, 1997). Initial ideas of encoding variability were supported by experiments demonstrating the positive effects of spaced as opposed to massed learning, suggesting that longer repetition lags lead to more independent representations of repeated stimuli (e.g., Bray, Robbins, & Witcher, 1976). However, not all studies reported encoding variability to have beneficial effects on memory performance (see Postman & Knecht, 1983). Although, generally, repetition appears to improve subsequent recognition memory, it was also suggested that it reduced our ability to correctly reject similar lures (Reagh & Yassa, 2014). Those findings supported the notion that repetition enhances item memory but at the cost of contextual details, as proposed by the competition trace theory (Yassa & Reagh, 2013). It appears therefore, that repeated encoding of a stimulus may have dissociative effects on subsequent item and source memory (e.g., context) performance (see also Sievers & Renoult, 2019).

Results from retroactive interference paradigms are typically in line with the competition trace theory. Retroactive interference can be measured using an AB/AC paradigm (Postman & Underwood, 1973), where stimulus A is first encountered in context B, followed by another presentation of stimulus A in another, interfering context C. Studies employing this paradigm have consistently reported inferior context memory compared to presenting a stimulus just once, in a single context (Anderson & Neely, 1996; Hupbach, Gomez, Hardt, & Nadel, 2007; Kim et al., 2017; McGovern, 1964). It is thought that repeating the stimulus in a novel context reactivates the memory associated with the first context (McClelland, McNaughton, & O’Reilly, 1995; Norman & O’Reilly, 2003), which is in line with predictions based on the reactivation view. The new context, C, is then integrated in an effort to generalise across contexts B and C, weakening subsequent source memory (Richter, Chanales, & Kuhl, 2016; Schlichting & Preston, 2015; Schlichting, Zeithamova, & Preston, 2014; Shohamy & Wagner, 2008; Zeithamova & Preston, 2010). Analogously, the competition trace theory suggests that the competition between those non-overlapping contextual features results in poorer context memory (Yassa & Reagh, 2013). This is in contrast to the multiple trace theory, which postulates that each stimulus encounter is encoded uniquely, enhancing subsequent retrieval of episodic details (Hintzman, 1986; Hintzman & Block, 1971; Nadel & Moscovitch, 1997). It is noteworthy that retroactive interference paradigms only present stimuli twice and do not consider possible encoding variability effects after more than two presentations. Moreover, retroactive interference may explain poorer context memory, but not much attention has been paid to possible mechanisms supporting correct context memory in retroactive interference or encoding variability paradigms. It may be that reactivation and the creation of multiple traces contribute to different aspects of memory formation when information is encoded in varying contexts. For example, Opitz (2010) reported that encoding stimuli in different contexts strengthened item memory but the memory judgements relied more on familiarity than episodic recollection.

Reactivation of neural patterns relating to memory encoding can be tested with representational similarity analysis (RSA; Kriegeskorte, Mur, & Bandettini, 2008). Pattern reactivation is indexed by a correlation coefficient, which is computed between repetitions of stimuli across voxels. Results from functional magnetic resonance imaging (fMRI) research have generally supported the reactivation view by reporting higher pattern similarity between repeated encoding presentations for subsequently remembered compared to forgotten stimuli (e.g., Levy Wagner, 2013; Staresina, Alink, Kriegeskorte, & Henson, 2013; van den Honert, McCarthy, & Johnson, 2016; Xue et al., 2010). The majority of those studies repeatedly presented participants with the same stimuli under the same encoding instructions before testing their memory, and thus the use of the same encoding task repeatedly may have potentially favoured reactivation. Studies employing retroactive interference paradigms have provided only partial support for the reactivation view. For example, pattern similarity during retroactive interference suggested that reactivation predicted successful source memory but only when interference was low, i.e., when context C was forgotten (Koen & Rugg, 2016). This was observed in task-selective voxels for the four different tasks. Contrary to predictions based on the reactivation framework, another study reported a negative relationship in the lateral occipital cortex between subsequent source memory and pattern reactivation between item repetitions (Kim, Norman, Turk-Browne, 2018). The authors suggested that reactivation of item-specific patterns explained the phenomenon of retroactive interference. Although not discussed, those results may indicate that another process, such as the creation of multiple traces, is related to the formation of correct context memory, i.e., higher resistance to context interference. Taken together, results from these retroactive interference paradigms may seem somewhat incompatible, however, it is conceivable that different brain regions perform distinct processes. It may be that higher reactivation in task-related regions, as functionally defined by Koen and Rugg (2016), contributes to source memory formation, while reactivation in the lateral occipital cortex signals generalisation across tasks. Interestingly, representational variability in the superior parietal cortex was shown to be associated with superior long-term memory retention in a recurrent-testing paradigm, i.e., when memory is explicitly tested at each presentation of the stimulus (Karlsson Wirebring et al., 2015), adding support for the encoding variability view. However, none of these previous studies directly compared similarity patterns relating to correct and incorrect context memory. Additionally, AB/AC paradigms do not contrast repeated encoding in the same or multiple, varying contexts but instead investigate pattern similarity for different memory outcomes only during encoding variability. Moreover, these studies compare behavioural memory performance at a single presentation and multiple presentations in different contexts, which constitutes an unfair comparison because repetition improves memory performance.

Research has investigated the effect of encoding variability on subsequent memory performance, mainly in retroactive interference paradigms, but very few studies have directly compared the behavioural and neural differences between repeatedly performing the same task and performing different tasks, i.e., encoding repeated stimuli in the same context or different contexts. Such a paradigm allows a direct comparison between the reactivation hypothesis (Hintzman, 2004, 2010) and encoding variability view (Bower, 1972; Hintzman, 1986). In the present fMRI experiment, participants encoded stimuli (famous names) under two encoding context conditions: 1) encode stimuli four times in the same context, i.e., same judgement task, 2) encode stimuli four times in different contexts, i.e., different judgement tasks at each presentation. Following the encoding phase, item memory (famous names) and source memory (context) were tested. Participants encoded one-half of a set of stimuli in the same context, hypothesised to be supported by pattern reactivation as proposed by the reactivation view. The other half of the stimuli was encoded in different contexts, which is hypothesised to involve one of two different operations, reactivation or the creation of multiple traces. We predict reactivation, as reflected in higher pattern similarity, to be associated with incorrect context judgements. This is in line with results from retroactive interference paradigms, suggesting that reactivation indexes higher levels of generalisation across contextual information (Richter et al., 2016; Schlichting & Preston, 2015; Schlichting et al., 2014; Shohamy & Wagner, 2008; Zeithamova & Preston, 2010). Subsequent correct context memory judgements, on the other hand, are predicted to be associated with lower levels of pattern reactivation, possibly reflecting the creation of multiple traces, as proposed by the encoding variability view and multiple trace theory (Hintzman, 1986; Hintzman Block, 1971; Nadel & Moscovitch, 1997). Therefore, subsequent source memory performance can be predicted based on the cognitive and neural operations involved during different context encoding.

## Materials and Methods

### Overview

This study employed a subsequent recognition-source memory paradigm with a study and a test phase. The study phase was split into four blocks during which participants were presented with written names of famous people. Each stimulus was presented four times. Half of the stimuli were repeatedly encoded in the same context, i.e., participants answered the same question about the famous person four times (e.g., “Is this person female?”). The other half of the stimuli were presented in different contexts, i.e., participants answered a different question at each of the four presentations of the stimulus (e.g., “Is this person female?”, “Is this person currently active in show business?”, “Is this person British?” and “Do you like this person?”). The study phase was followed by a short break before the test phase began. During the test phase, participants were presented with all the names from the study phase, which were interspersed with other famous names they had not seen before in the experiment. Participants were instructed to make an old/new judgement, including a confidence rating. Whenever a stimulus was rated as old, i.e., presented during the study phase, participants were presented with a source memory recognition task asking in which context the stimulus had previously been encountered; one of the judgement tasks or all of them. This follow-up question assessed source memory. In the following, correct item and source memory judgements are termed *hits*^+^, correct item but incorrect source memory judgements are termed *hits*^−^ and incorrect item memory judgements are labelled *misses*.

### Participants

Twenty British healthy adult volunteers (12 females) were recruited through opportunity sampling including mailing lists and posters. Participants were 18 to 36 years old (M_*age*_ = 24± 6), native English speakers, right-handed, with an average of 16 ± 3 years of education, normal or corrected-to-normal vision and confirmed they had never been diagnosed with any psychiatric or neurological disorder. Informed consent was obtained and this research was approved by the Research Ethics Board of the University of East Anglia. Participants were reimbursed for their time. Data from one participant were excluded from all analyses due to technical faults during scanning. Data from another two participants were excluded from imaging analyses due to excessive movement during the scan, resulting in *N* = 17 (10 females).

### Materials

Experimental tasks were programmed and delivered through the stimulus presentation software Presentation 18.1 (https://www.neurobs.com/). Stimuli were 240 written names of famous people (e.g., Keith Richards, Michelle Obama). Stimuli were selected from a total of 350 famous names based on a behavioural pilot study to identify the most well-known famous people amongst a sample of participants with similar characteristics as the sample in the present experiment. Famous names were matched in accordance with the four encoding tasks (gender, currently active in show business or not, British or not) across tasks and conditions. Stimuli were presented as written words in white Courier New 36 font, in the centre of a black background.

### Task & Procedure

The study phase was split across four blocks/runs. At the beginning of each block, participants were given a question to answer about the famous names they were going to see in the block. The four questions were “Is this person female?”, “Is this person currently active in show business?”, “Is this person British?” and “Do you like this person?”; all requiring yes/no answers by pressing one of two buttons. Task order was pseudo-randomised across participants. When participants did not know the answer or were unfamiliar with the famous name, they were encouraged to guess the answer. Half of the stimuli were presented repeatedly within only one of the tasks (*same context*), the other half was presented once in each of the four tasks (*different context*). This resulted in a total of 480 encoding trials, with an average inter-trial interval (ITI) of 4100 ms. The experimental procedure and trial timings are illustrated in Figure 1. At the end of the study phase, participants had a short break during which they could rest their eyes.

**Figure 1.**
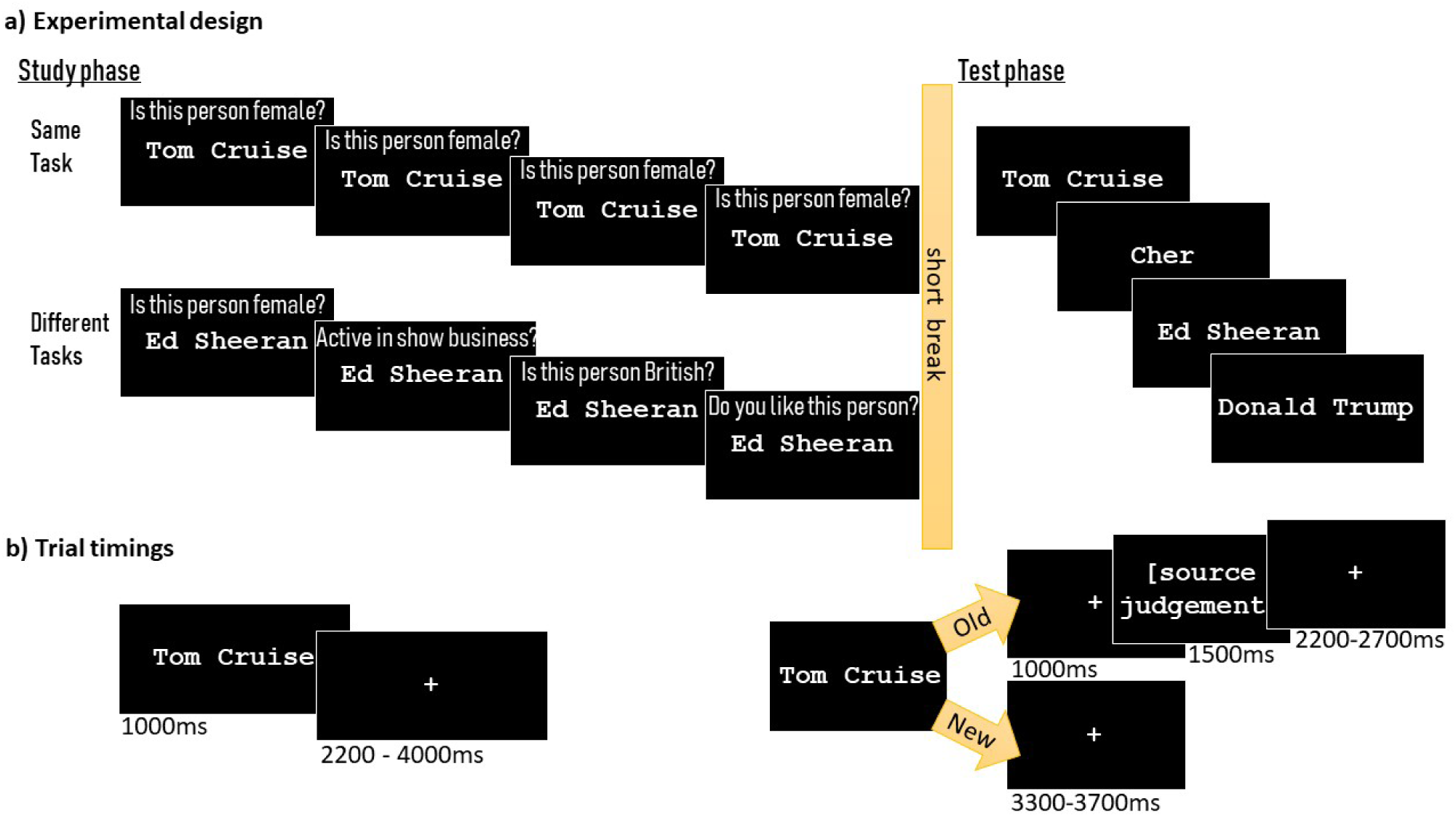
a) The experimental paradigm with four encoding presentations of each stimulus during the study phase; participants encoded half of the stimuli in a different task condition, i.e., performing a different task at each presentation of the stimulus, the other half were encoded in a same task condition, i.e., participants repeatedly performed the same encoding task; in a test phase, participants made old/new judgements followed by source judgements. *Note:* Task instructions/questions were presented at the beginning of each run rather than during each stimulus presentation as illustrated here; b) trial timings.

In the test phase, participants performed an unexpected recognition-source memory test. Stimuli from the encoding phase were presented along with the remaining new, i.e., previously unseen, stimuli. Both lists were matched in accordance with the four encoding tasks to have equal numbers of males/females, currently active/not active in show business and British/non-British names in either set. Participants gave an old/new response depending on whether or not they thought the name had been presented during the study phase by pressing one of four buttons on the response pad corresponding to the following responses: “definitely old”, “perhaps old”, “perhaps new”, and “definitely new”. “Old” responses were followed up with a source memory question asking participants which task they had previously performed for the stimulus with the response options “all four tasks”, “gender task”, “show business task”, “British task”, “like task” and “I don’t know”. Famous names were presented for 1500 ms, followed by a fixation cross for 1000 ms. Depending on the old/new response, either a fixation cross or the source memory question appeared for 1500 ms (see Figure 1b). Another fixation cross of random duration (800 – 1200 ms, average ITI = 5000 ms) indicated the beginning of the next trial.

### Imaging

Functional and anatomical MRI data were obtained with a 3 Tesla wide bore GE 750w MRI scanner. Stimuli and task instructions were presented on a screen in the scanner, approximately 90 cm from participants’ eyes, via an AVOTEC silent vision projector. Behavioural responses to the task were recorded with a Fiber Optic Response Device and logged with Presentation 18.1. Participants wore earplugs during the scan, head motion was reduced with foam pads. The structural T1-weighted image was acquired using GE’s gradient echo pulse sequence, BRAVO. Slices were angled parallel to the falx cerebri, the centre of the field of view (FOV) was angled to AC-PC (TR = 7.25, TE = 2.55, flip angle = 9°, FOV = 230 mm^2^, matrix size = 256 × 256, slice thickness = 0.9 mm, number of slices = 196). Functional 2T*-weighted images were acquired with a gradient-echo, EPI sequence (TR = 2000 ms, TE = 28 ms, flip angle = 90°, FOV = 213 mm^2^, matrix size = 64 × 64, slice thickness = 3 mm, number of slices per volume = 35, number of volumes = 260 for encoding blocks, 318 for recognition blocks). When full brain coverage was not possible, the FOV was angled to primarily include the temporal and parietal lobes as well as prefrontal cortex.

Pre-processing and analyses of imaging data were carried out in SPM12 (Wellcome Trust Center, London, UK, www.fil.ion.ucl.ac.uk) and Matlab (The Mathworks, Inc., Natick, MA, USA). The first six volumes of each run were discarded. Functional images were slice-timing corrected before spatial realignment because of the interleaved acquisition (Ashburner et al., 2016). Spatial realignment parameters were kept at default, 4 mm sampling distance, spatial smoothing with a 5 mm full-width-half-maximum (FWHM) Gaussian kernel, images were registered to their mean, 6th degree B-Spline interpolation method, no wrapping, no differential weighting of voxels. The structural image was coregistered to the mean functional image and then segmented into grey and white matter, bias corrected and spatially normalised to the Montreal Neurological Institute (MNI) space selecting forward deformations. The deformation parameters were used to spatially normalise functional images and the bias corrected structural image. Functional images were resampled to an isotropic voxels size of 3 mm, the structural image was resampled to 1 mm^3^ voxel size, both closely matched the original voxel sizes.

Representational similarity scores were calculated from the preprocessed, unsmoothed images. Customised explicit within-brain masks were used during model estimation. Single-trial betas were estimated with the *Least Squares-Separate* (LSS) approach (for more information see Mumford, Turner, Ashby, & Poldrack, 2012) and submitted to a whole-brain searchlight RSA (Kriegeskorte et al., 2008). For each trial, beta values were extracted for 5 × 5 × 5 voxel cubes centred around each voxel (see Wing, Ritchey, & Cabeza, 2015; similar results were obtained with 3 × 3 × 3 voxel cubes). Similarity scores were computed between stimulus presentations by correlating the winsorised (*SD* = 3) betas for each stimulus and each presentation pair, i.e., Presentations 1 & 2, Presentations 1 & 3, Presentations 1 & 4, Presentations 2 & 3, Presentations 2 & 4 and Presentations 3 & 4. The resulting six encoding similarity indices were Fisher transformed and then averaged. The images, which contained the 5 × 5 × 5 voxel-wise similarity scores were smoothed using a 6 mm^3^ FWHM Gaussian kernel before 2^*nd*^-level whole brain analyses were carried out in SPM12. To evaluate the relative contributions of pattern reactivation relating to individual study items and those associated with more task-general processes (see Wing et al., 2015), item-level and set-level similarity were calculated, illustrated in Figure 2. Item-level similarity was computed by correlating the beta values of a stimulus at one presentation with another presentation of the same stimulus. This measure reflects the degree to which stimulus properties are reactivated. Set-level similarity was calculated by correlating beta series for a specific stimulus at one presentation with all other stimuli in the same category (e.g., same task, source hits) at another presentation. The resulting correlation coefficients were averaged at the voxel level. This condition-wise between-item similarity provides an index of task-general information or processes that are shared between stimuli of the same category.

**Figure 2.**
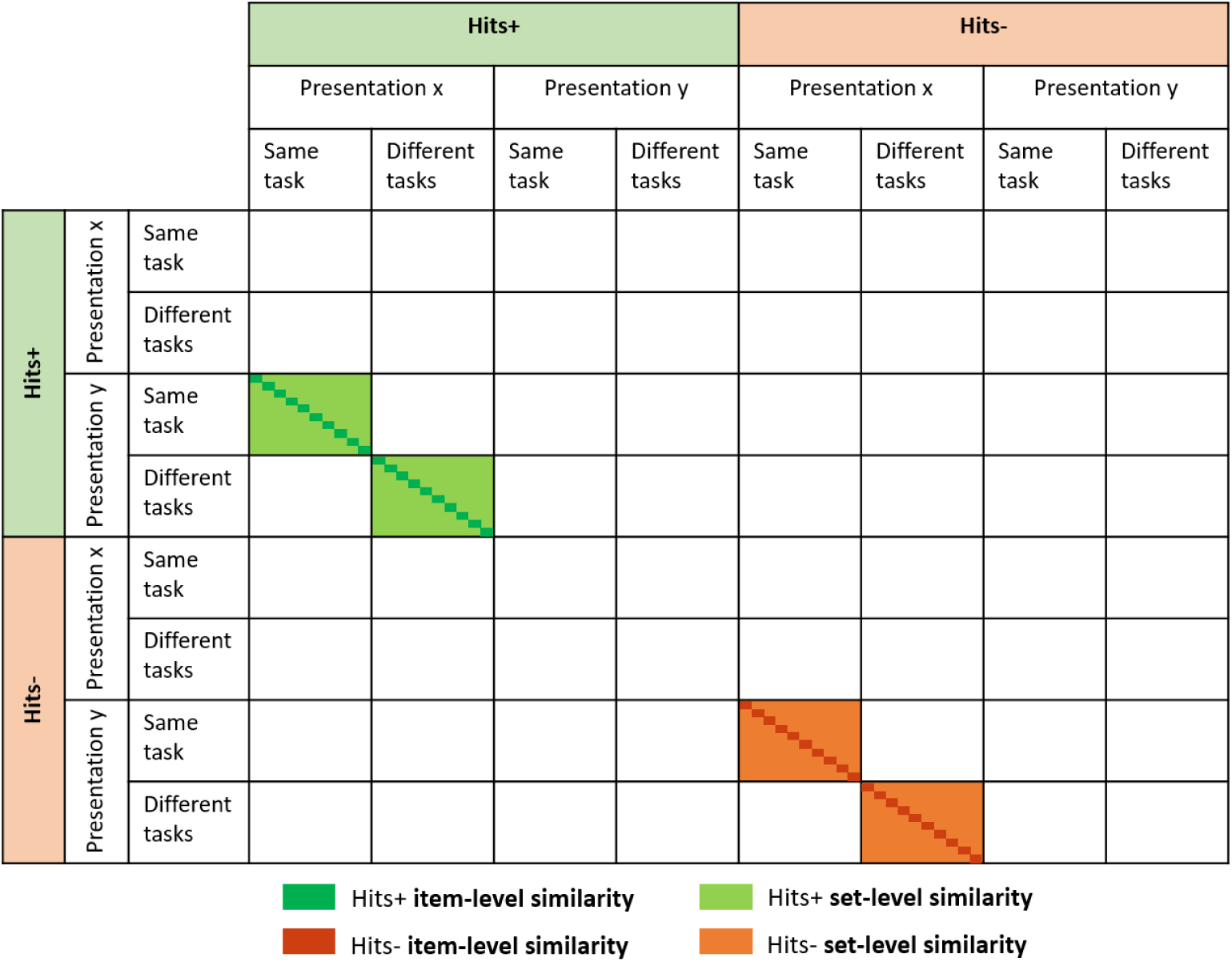
Identity matrix displaying the four conditions for which item- and set-level similarity scores were calculated.

Because same task repetitions occurred within the same fMRI runs, whereas different task repetitions occurred between runs, encoding similarity patterns for same and different task encoding could not be contrasted statistically, due to intrinsic variability in the fMRI signal between runs. However, this issue does not affect possible differences that might be observed in pattern similarity due to source memory performance within the encoding task conditions or the interaction between source memory performance and encoding task condition. Such an interaction directly tests the hypothesis that in the same task condition, hits^+^ judgements are associated with more reactivation than hits^−^ judgements; while in the different task condition, hits^−^ judgements are associated with higher levels of reactivation than hits^+^ judgements.

## Results

### Behavioural results

#### Discriminability analysis

The normalised probabilities of overall hits and false alarms were compared in a paired-samples *t*-test. The *t*-test revealed that participants’ item memory performance was above chance, *t*_16_ = 17.180, *p* ≤ .001. Mean and standard deviations of discriminability scores, *d*’, and percentages of hits and false alarms are displayed in Table 1. Those individual *d*’ scores indicate that item recognition memory performance was higher in the different encoding task condition, however, this difference was statistically non-significant, t_16_ = 0.334, *p* = .743.

**Table 1.** The mean d’ scores and mean % of Hits and False Alarms with standard deviations (in brackets) for overall memory performance and across the two encoding conditions, different and same task encoding.

| | $d'$ (SD) | $M_{\text{Hits}}\%$ (SD) | $M_{\text{FalseAlarms}}\%$ (SD) |
| --- | --- | --- | --- |
| Overall | 2.79 (0.69) | 85.59 (11.25) | 7.21 (5.16) |
| Different task | 3.22 (1.37) | 89.51 (9.62) | 7.21 (5.16) |
| Same task | 3.06 (1.72) | 81.67 (13.60) | 7.21 (5.16) |

#### Behavioural performance at test

Frequencies of hits^+^, hits^−^ and misses (memory performance) were analysed in relation to encoding task condition (same, different) with a 3 × 2 repeated-measures ANOVA. Descriptive statistics are displayed in Figure 3. The ANOVA revealed an interaction between the two factors, memory performance and encoding context, *F*_1,21_ = 5.335, *p* = .027. A post-hoc test following up on the interaction showed that encoding under the different encoding task condition was associated with more hits^−^ judgements (*M*_%_ = 62.02, *SD* = 17.25) than the same encoding task condition (*M*_%_ = 48.68, *SD* = 12.00), *p* = .013. The same encoding task condition was associated with more misses (*M*_%_ = 18.51, *SD* = 13.13) than the different encoding task condition (*M*_%_ = 10.70, *SD* = 9.60), *p* ≤ .001. Finally, as illustrated in Figure 3, there were more hits^+^ judgements in the same (*M*_%_ = 32.81, *SD* = 17.53) than in the different task condition (*M*_%_ = 27.28, *SD* = 19.15), but this was statistically non-significant, *p* = 0.281.

**Figure 3.**
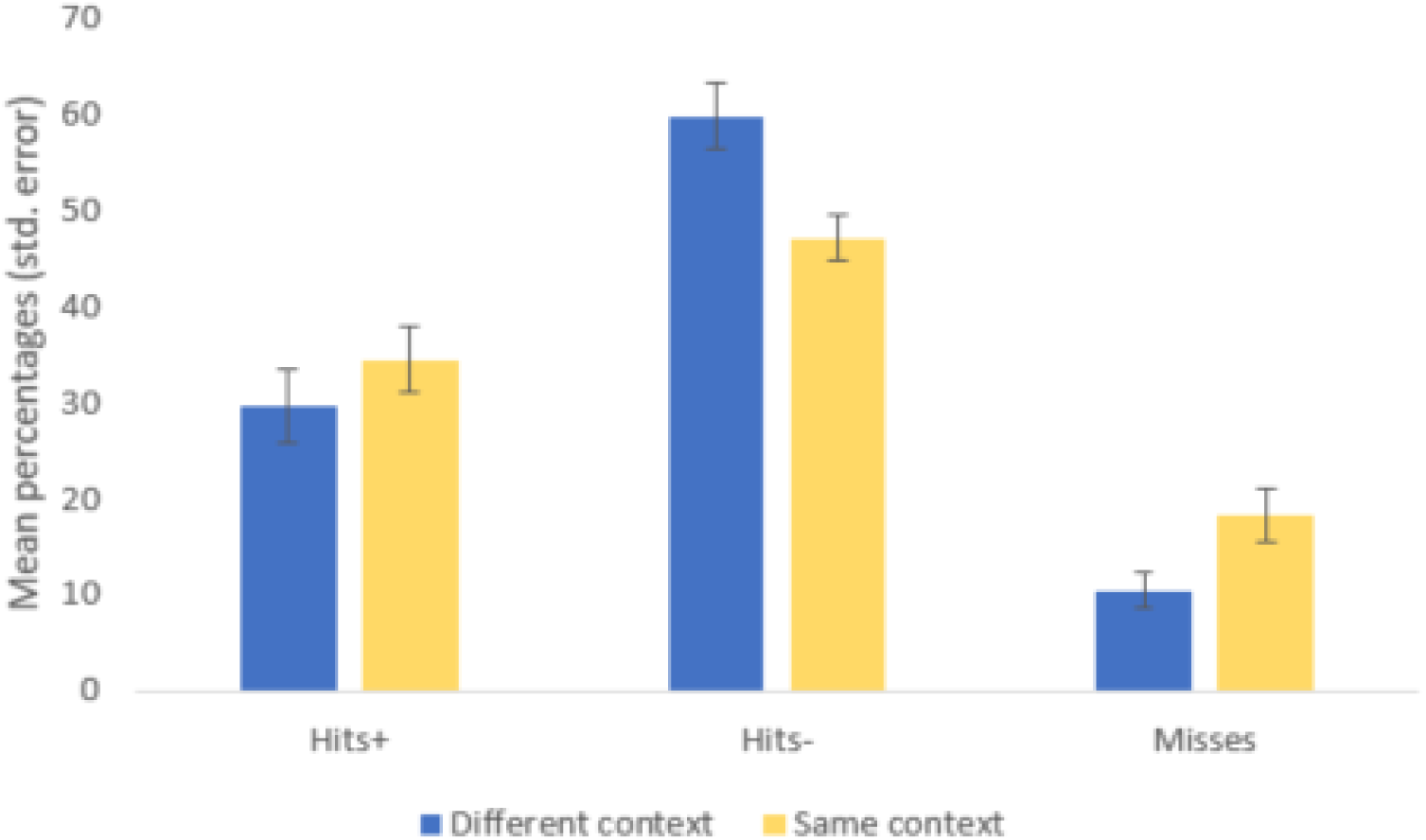
The mean percentages and standard errors of the three levels of memory performance (hits^+^, hits^−^, misses) and encoding context (different, same).

### Representational similarity analysis

#### Searchlight analysis

Pattern similarity was calculated across repeated study episodes between all possible encoding presentation pairs, i.e., Presentations 1 & 2, Presentations 1 & 3, Presentations 1 & 4, Presentations 2 & 3, Presentations 2 & 4 and Presentations 3 & 4. This resulted in six encoding similarity indices that were averaged. Due to low frequencies of misses, the neuroimaging analyses focused on trials resulting in hits^+^ and hits^−^ judgements. Separate whole-brain encoding similarity indices were computed for each of the four conditions, Hits^+^ Same Context, Hits^−^ Same Context, Hits^+^ Different Context, Hits^−^ Different Context. The hypothesised interaction between source memory and encoding context was tested with a *t*-contrast (contrast weights: 1, −1,-1, 1, *p* ≤ .001), modelling pattern activation to be higher for Hits^+^ Same Context and Hits^−^ Different Context than for Hits^−^ Same Context and Hits^+^ Different Context. As illustrated in Figure 4, the whole-brain analysis revealed one statistically significant cluster located in the posterior cingulate cortex (*k* = 175, *p* = .030, *FWE*-corrected; peak voxel MNI_*x,y,z*_ = −21, −40, 29, *t* = 4.57, *p* = 0.041, *FWE*-corrected).

**Figure 4.**
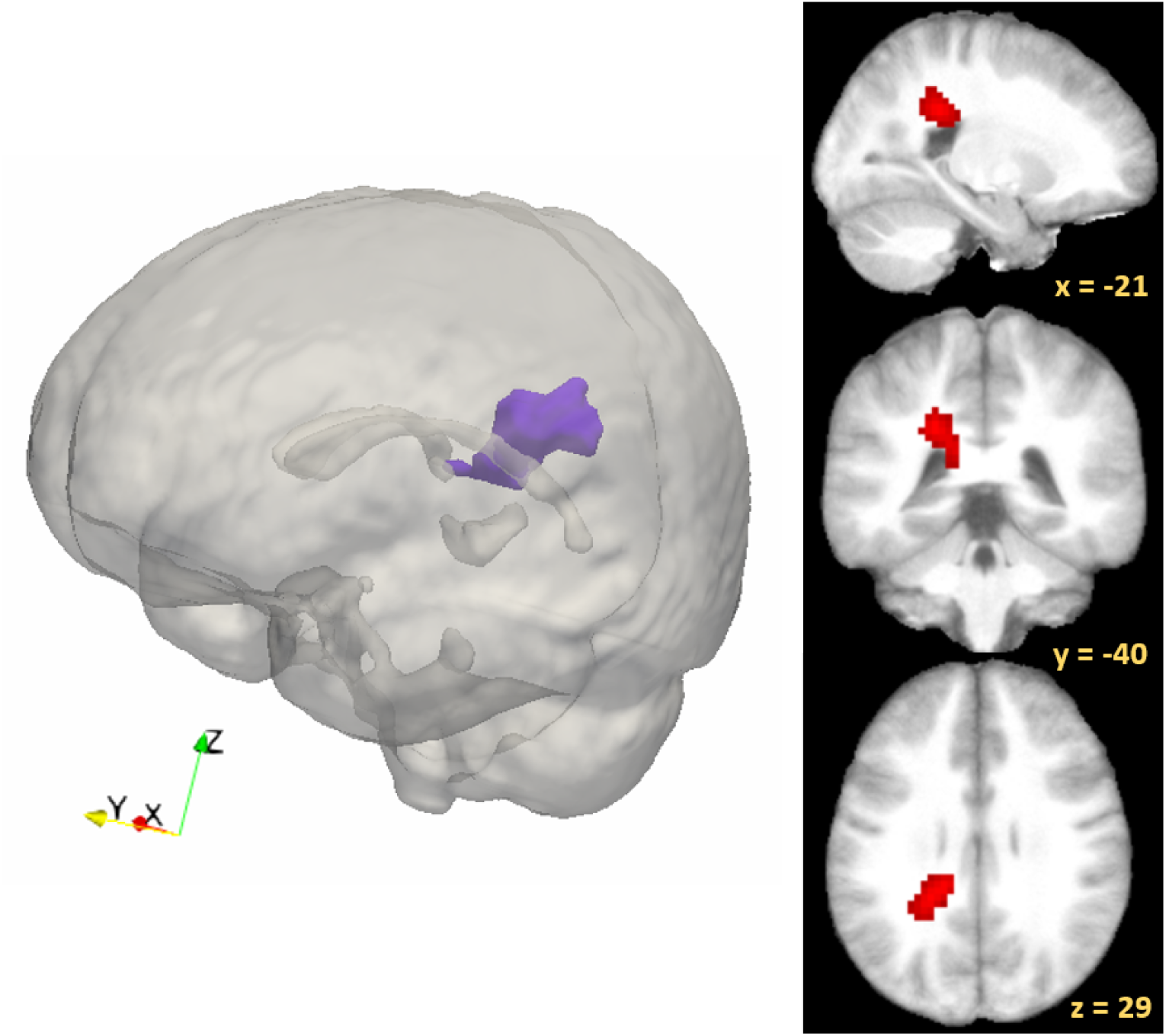
The predicted interaction between source memory and encoding task condition as observed in a cluster of voxels located in the posterior cingulate cortex. The glassbrain illustration was constructed using ITK-SNAP (Yushkevich et al., 2006) and ParaView (Ahrens, Geveci, & Law, 2005; described in Madan, 2015). The brain slices were obtained in SPM.

#### Item-level and set-level similarity

To evaluate the relative contributions of item- and set-level similarity, a mask was created based on the 175 voxels in the significant cluster revealed by the whole-brain analysis (see Figure 4). Indices were calculated for item- and set-level similarity for the four conditions of interest, and separate 2 × 2 repeated-measures ANOVAs were performed for item-level and set-level similarity, each with the factors subsequent source memory (Hits^+^, Hits^−^) and encoding context (same task, different tasks). Mean similarity scores are displayed in Figure 5. The ANOVA on item-level similarity scores revealed a strong, but statistically non-significant trend for the interaction, *F*_1,16_ = 3.519, *p* = .079. For set-level similarity, the interaction between the two factors was significant, *F*_1,16_ = 32.487, *p* ≤ .001. The interaction is clearly illustrated in the bar graph in Figure 5, however, appears to be similar for item- and set-level similarity. To confirm that there were no differences in the interaction between item-level and set-level similarity, a further 2 × 2 × 2 ANOVA was performed, including similarity type as a factor. The 3-way interaction between source memory, encoding task condition and similarity type was not significant, *F*_1,16_ = 0.104, p = .751.

**Figure 5.**
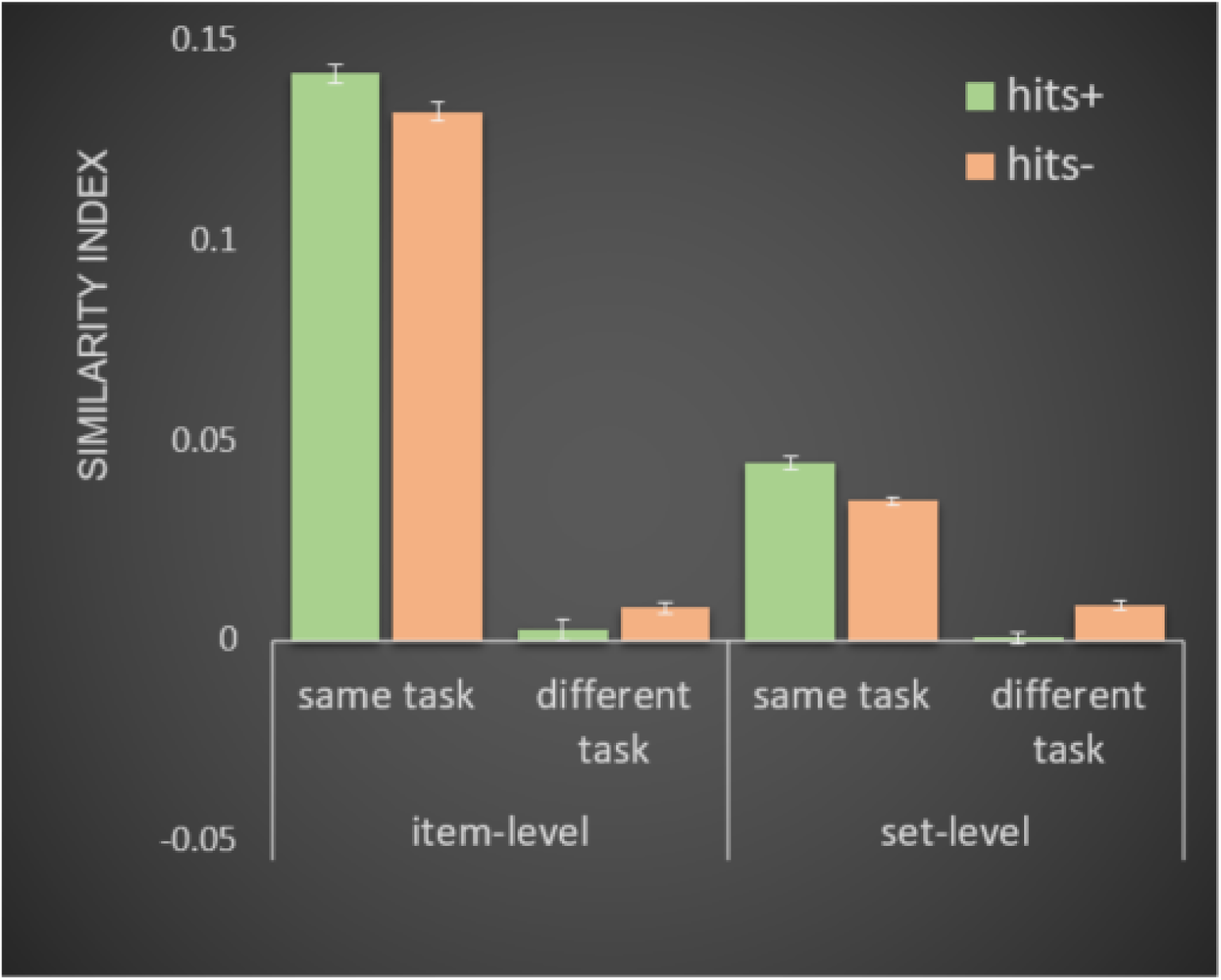
Item- and set-level similarity scores in posterior cingulate cortex cluster for subsequent hits^+^ and hits^−^ judgements in the two encoding conditions, same and different task encoding. Error bars denote standard error.

## Discussion

The present experiment investigated how neural pattern reactivation supports successful memory formation across multiple study episodes. We tested for differences in neural similarity patterns relating to source memory effects when stimuli were either repeated in the same context or repeatedly encoded in differing contexts. Behavioural results showed that different task encoding, when compared to same task encoding, was associated with better item memory but poorer source memory performance. Neural pattern similarity was assessed using representational similarity analysis. The results supported the hypothesis that dissociable neural mechanisms underlie source memory formation in same and different context encoding. The predicted interaction between encoding task condition and source memory performance was observed in the posterior cingulate cortex. Higher levels of pattern reactivation are associated with superior source memory performance when stimuli are repeatedly encoded in the same context. In the different encoding task condition, pattern reactivation was lower for subsequent correct item and source memory judgements (hits^+^) than for correct item but incorrect source memory judgements (hits^−^). These findings suggest that the creation of multiple memory traces, rather than reactivation of existing traces, supports the successful encoding of context memory during encoding variability.

Behavioural results supported the hypothesis that the different task condition was associated with fewer misses, i.e., superior item memory, but also with more hits-judgements, i.e., worse source memory, than the same encoding task condition. These results largely replicate recent behavioural data from our lab using the same paradigm (Sievers & Renoult, 2019; see also Opitz, 2010; Reagh & Yassa, 2014). The reported interaction between memory performance and encoding task condition is reminiscent of results from retroactive interference paradigms (Anderson & Neely, 1996; Hupbach et al., 2007; Kim et al., 2017). This line of research has indicated that stimulus occurrence in multiple contexts may cause interference, resulting in higher levels of generalisation at the cost of contextual source information. Despite weaker source memory performance, participants were less likely to forget the stimulus itself, when it was encoded in differing contexts, suggesting that encoding variability is associated with better item memory (Bower, 1972; Hintzman, 1986; Nadel & Moscovitch, 1997). Overall, the behavioural results provide additional support for the notion that encoding variability is associated with stronger item memory but worse performance in a source memory task requiring the retrieval of varying encoding contexts (Sievers & Renoult, 2019).

Neural similarity patterns across multiple encoding episodes supported the hypothesis that pattern reactivation differentially predicted source memory performance depending on the encoding task condition. Reactivation, as indexed by higher pattern similarity, was beneficial for source memory when items were repeatedly encoded in the same context. However, when items were repeatedly encoded in different contexts, reactivation was predicted to facilitate generalisation of contextual information, which would result in poorer source memory for the multiple contexts the items were initially encoded in. A searchlight analysis revealed the hypothesised interaction in a set of voxels located in the left posterior cingulate cortex. In the same task condition, pattern reactivation in this brain region was higher for subsequent hits^+^ than hits^−^ judgements, with the opposite pattern observed in the different task condition. For the same task condition, this result is consistent with a large body of research supporting the reactivation view (Benjamin & Tullis, 2010; Thios & D’Agostino, 1976), which posits that reactivation of the same patterns facilitates successful memory encoding (van den Honert et al., 2016; Ward, Chun, & Kuhl, 2013; Xue et al., 2010). For the different task condition, this interaction suggests that another process, potentially the formation of multiple non-identical traces, supports the successful formation of multiple context memory (Bower, 1972; Hintzman, 1986; Martin, 1968; Nadel & Moscovitch, 1997). Posterior parietal regions, including the posterior cingulate cortex and precuneus, have repeatedly been implicated in memory encoding and consolidation (see Gilmore, Nelson, & McDermott, 2015; Rugg & King, 2017; Sestieri, Shulman, & Corbetta, 2017). Moreover, a meta-analysis reported the posterior cingulate gyrus to be one of seven regions that make up a semantic network in the brain (Binder, Desai, Graves, & Conant, 2009). The authors noted that this posterior parietal region was typically associated with episodic memory processes, however, it may act as the intersection between semantic retrieval and episodic encoding. This is in line with suggestions that left parietal regions are a part of a larger network that is involved during processes of semantic feature integration (Binder et al., 2009; Chou, Chen, Wu, & Booth, 2009; Fairhall & Caramazza, 2013). Existing research has also shown that pattern similarity in the posterior cingulate cortex, as computed between encoding and rehearsal of video clips, was related to the amount of details that could later on be recalled (Bird, Keidel, Ing, Horner, & Burgess, 2015). The authors suggested that reinstatement in those posterior midline structures facilitated consolidation of complex events, which become more generic and somewhat less episodic through this consolidation process. The present results extend this hypothesis by demonstrating that different context encoding is associated with lower pattern similarity in order to preserve unique contextual details.

Following up on the searchlight results, two measures of similarity were computed for the significant cluster: item-level and set-level similarity. Item-level similarity is thought to reflect reactivations of stimulus properties, while set-level similarity indexes reactivations of more general processes and information that is shared between all stimuli within a particular category (e.g., Wing et al., 2015), e.g., different task, hits^+^ judgements. The predicted interaction, reflecting higher pattern reactivation to be associated with better subsequent context memory in the same task condition but worse context memory in the different task condition, was observed in both item- and set-level similarity (see Figure 5), though only statistically significant for set-level similarity. Moreover, the interaction did not differ between item- and set-level similarity. This indicates that we not only observe the dissociation in pattern reactivation with respect to source memory performance between same and different task encoding, but also at a more general set-level. For the same encoding task condition, the set-level result suggests that items for which the source can subsequently be retrieved are more similarly represented to each other, when compared to items for which the source cannot be retrieved later on. This suggests that processes and features, which are shared between subsequent hits^+^ judgements, are reactivated during same context repetitions. In the different task condition, items resulting in subsequent hits^+^ judgements were less similar to each other than subsequent hits^−^ judgements. These findings support the prediction that higher levels of generalisation were associated with worse subsequent source memory performance. It further indicates that the processes underlying successful encoding of different contexts were more different between items, potentially providing support for the encoding variability view and the multiple trace theory (Bower, 1972; Hintzman, 1986; Martin, 1968; Nadel & Moscovitch, 1997). Taken together, these findings are consistent with previous research, showing that reactivation of shared item features, possibly indexing generalisation, was associated with worse source memory for the encoding task (Kim et al., 2018).

In conclusion, the present results provide support for the notion that same and different context encoding are supported by distinct mechanisms. Mere repetition of an item in the same context strengthens item and source memory by reactivation. When items are repeatedly encoded in different contexts, reactivation appears to promote generalisation at the cost of contextual features. Another mechanism, possibly the creation of multiple distinct memory traces, underlies the successful encoding of multiple contexts. The behavioural results indicated that different context encoding was associated with better item, but worse source memory, than same context encoding. Previous research had largely focused on the mechanisms underlying the forgetting of episodic details, with less emphasis on the mechanisms underlying successful context encoding. It is noteworthy that, although generalisation in the context of different task encoding was associated with worse source memory, such processes of abstraction and generalisation can often be very useful, as it has been suggested to be the basis of semantic knowledge (e.g., Binder & Desai, 2011; Cermak, 1984). In other words, while contextual information is sometimes important to be remembered, the creation of a more coherent, semantic memory representation is critical for making memory-guided decisions and future inferences.

## Acknowledgements

This work was supported by a grant number 132/14 from the BIALfoundation [“How memories form”] to Louis Renoult and Fraser W Smith.

